# An optimized CRISPR/Cas toolbox for efficient germline and somatic genome engineering in *Drosophila*

**DOI:** 10.1101/003541

**Authors:** Fillip Port, Hui-Min Chen, Tzumin Lee, Simon L. Bullock

**Affiliations:** MRC-Laboratory of Molecular Biology, Francis Crick Avenue, Cambridge CB2 0QH, UK; Howard Hughes Medical Institute, Janelia Farm Research Campus, Ashburn, Virginia VA 20147, USA

## Abstract

The type II CRISPR/Cas system has recently emerged as a powerful method to manipulate the genomes of various organisms. Here, we report a novel toolbox for high efficiency genome engineering of *Drosophila melanogaster* consisting of transgenic Cas9 lines and versatile guide RNA (gRNA) expression plasmids. Systematic evaluation reveals Cas9 lines with ubiquitous or germline restricted patterns of activity. We also demonstrate differential activity of the same gRNA expressed from different U6 snRNA promoters, with the previously untested U6:3 promoter giving the most potent effect. Choosing an appropriate combination of Cas9 and gRNA allows targeting of essential and non-essential genes with transmission rates ranging from 25% - 100%. We also provide evidence that our optimized CRISPR/Cas tools can be used for offset nicking-based mutagenesis and, in combination with oligonucleotide donors, to precisely edit the genome by homologous recombination with efficiencies that do not require the use of visible markers. Lastly, we demonstrate a novel application of CRISPR/Cas-mediated technology in revealing loss-of-function phenotypes in somatic cells following efficient biallelic targeting by Cas9 expressed in a ubiquitous or tissue-restricted manner. In summary, our CRISPR/Cas tools will facilitate the rapid evaluation of mutant phenotypes of specific genes and the precise modification of the genome with single nucleotide precision. Our results also pave the way for high throughput genetic screening with CRISPR/Cas.

## Introduction

Experimentally induced mutations in the genomes of model organisms have been the basis of much of our current understanding of biological mechanisms. However, traditional mutagenesis tools have significant drawbacks. Forward genetic approaches such as chemical mutagenesis lack specificity, leading to unwanted mutations at many sites in the genome. Traditional reverse genetic approaches, such as gene targeting by conventional homologous recombination, suffer from low efficiency and are therefore labor intensive. Novel methods have been developed in recent years that aim to modify genomes with high precision and high efficiency by introducing double stand breaks (DSBs) at defined loci (1). DSBs can be repaired by either non-homologous end joining (NHEJ) or homologous repair (HR). NHEJ is an error prone process that frequently leads to the generation of small, mutagenic insertions and deletions (Indels). HR repairs DSBs by precisely copying sequence from a donor template, allowing specific changes to be introduced into the genome (2).

The type II clustered regular interspersed short palindromic repeat (CRISPR)/Cas system has recently emerged as an extraordinarily powerful method for inducing site-specific DSBs in the genomes of a variety of organisms. The method exploits the RNA-guided endonuclease Cas9, which plays a key role in bacterial adaptive immune systems. Target specificity of Cas9 is encoded by a 20-nt spacer sequence in the crisprRNA, which pairs with the transactivating RNA to direct the endonuclease to the complementary target site in the DNA (3). For genome engineering, crisprRNA and transactivating RNA can be combined in a single chimeric guide RNA (gRNA), resulting in a simple two component system for creation of DSBs at defined sites (3). Binding of the Cas9/gRNA complex to a genomic target site is only constrained by the requirement for a short protospacer adjacent motif (PAM) next to the target site, which for the commonly used *Streptococcus pyogenes* Cas9 is NGG (4).

Several groups recently demonstrated CRISPR/Cas-mediated editing of the genome of *Drosophila melanogaster*, a key model organism for biological research (5–11). However, the rate of mutagenesis has varied widely both within and between different studies. Differences in the methods used to introduce Cas9 and gRNAs into the fly are likely to contribute significantly to different experimental outcomes. Kondo and Ueda expressed both Cas9 and gRNA from transgenes stably integrated into the genome (8), whereas all other studies have used microinjection of expression plasmids or *in vitro* transcribed RNA into embryos to deliver one or both CRISPR/Cas components (5–7, 9–11). Much of the currently available evidence suggests that transgenic provision of Cas9 increases rates of germline transmission substantially (8, 10, 11). However, the influence of different regulatory sequences within Cas9 transgenes on the rate of generating mutations, as well as their location within the organism, has not been evaluated. The effect of different promoter sequences on the activities of gRNAs has also not been explored systematically. It is therefore possible that suboptimal tools are currently being employed for many CRISPR/Cas experiments in *Drosophila*.

Previous studies in *Drosophila* have focused on the use of CRISPR/Cas to create heritable mutations in the germline. In principle, efficient biallelic targeting within somatic cells of *Drosophila* would represent a powerful system to dissect the functions of genes within an organismal context. However, the feasibility of such an approach has so far not been explored.

Here, we present a versatile CRISPR/Cas toolbox for *Drosophila* genome engineering consisting of a set of systematically evaluated transgenic Cas9 lines and gRNA expression plasmids. We describe combinations of Cas9 transgenes and *U6-gRNAs* that can be used to induce, with high efficiency, loss-of-function mutations in non-essential or essential genes and integration of short designer sequences by HR. Finally, we show that our optimized transgenic tools permit efficient biallelic targeting in a variety of somatic tissues of the fly, allowing the characterization of mutant phenotypes directly in Cas9/gRNA expressing animals.

## Results

### Generation and evaluation of Cas9 transgenes

We generated a series of Cas9-expressing transgenes in order to compare their expression patterns and endonuclease activities. Expression plasmids were produced encoding *S. pyogenes* Cas9 (codon optimized for expression in human (12, 13) or *Drosophila* cells) fused to nuclear localization signals under the control of different regulatory sequences. We generated two constructs in which Cas9 was under the control of the *nos* promoter, which is active in the male and female germline (14), and *nos* 3’UTR which targets protein synthesis to the germ cells (14). These constructs are similar to those used by two previous studies to provide Cas9 transgenically (8, 10). We also generated two novel constructs: *act-cas9* has a fusion of Cas9 to the ubiquitously expressed *actin5C* (*act*) promoter and the SV40 3’UTR and *UAS-cas9* has yeast *upstream activating sequences* (*UAS*), which allow expression under the control of the Gal4/UAS system (15), and the *p10* 3’UTR which promotes efficient translation (16). These plasmids were then integrated into the *Drosophila* genome at defined positions using the attB/attP/Phi31C system (17, 18). We also obtained two published stocks that express, from different genomic locations, a Cas9 transgene under control of the *vasa* promoter and 3’UTR, which is purported to restrict expression of the endonuclease to the germline (11). Details of Cas9 transgenes are provided in Table S1.

To functionally test the different Cas9 lines we designed a gRNA targeting the *ebony* (*e*) gene on the 3rd chromosome (referred to as *gRNA-e;* Fig. 1*A*). Mutation of both wild-type *e* alleles leads to very dark coloration of the adult cuticle. *gRNA-e* targets Cas9 to the 5’end of the *e* coding sequence, 25bp after the translation initiation codon. A construct expressing *gRNA-e* from the promoter of the U6:3 spliceosomal snRNA gene (see below) was stably integrated at the attP2 site on chromosome 3 (Table S2). Transgenic supply of the gRNA was designed to eliminate the variability between experiments that is associated with direct injection of a plasmid or *in vitro* transcribed RNA into embryos, thereby facilitating meaningful comparison of the activities of the different Cas9 lines.

**Figure 1:**
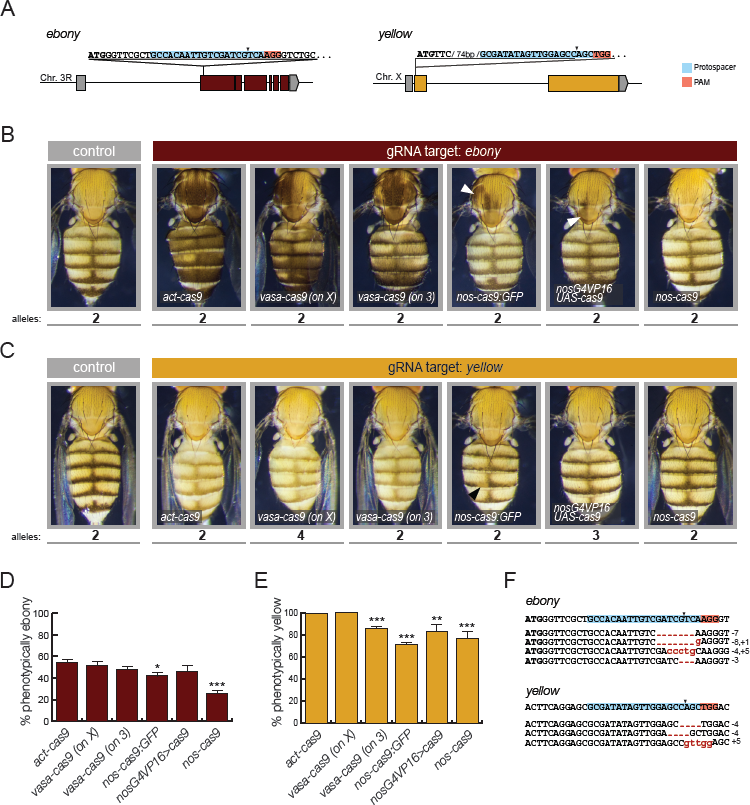
Systematic evaluation of transgenic Cas9 strains. (*A*) Schematic of sequence and position of *gRNA-y* and *gRNA-e* target sites in the *ebony* (*e*) and *yellow* (*y*) loci. Gray, untranslated regions. Both gRNAs target the 5’ end of the coding sequence (colored blocks) so that out-of-frame mutations should be loss-of-function alleles. Arrowheads, Cas9 cut site. (*B*, *C*) Representative examples of female flies expressing one copy of the different Cas9 transgenes and one copy of the *U6:3-gRNA-e* (*B*) or *U6:3-gRNA-y* (*C*) transgene compared to wild-type controls (at least 200 adults of each genotype were examined). In *B* and *C*, respectively, darker and lighter body coloration than the control indicates CRISPR/Cas mediated mutagenesis of the target gene in epidermal cells. The number of wild-type alleles present in each animal that have to be mutated to give rise to the mutant phenotype is indicated below each image. Somatic targeting is widespread in animals expressing *act-cas9* or *vasa-cas9*, sporadic in animals expressing *nos-cas9:GFP* and *nosG4VP16 UAS-cas9* (arrowheads) and undetectable in animals expressing *nos-cas9*. (*D*, *E*) Assessment of germline transmission of non-functional *e* (*D*) and *y* (*E*) mutant alleles induced by different Cas9 transgenes, performed by crossing animals expressing *cas9* and the appropriate gRNA to *e* or *y* mutant animals. Mean and SEM are shown from at least eight (*D*) or four (*E*) independent crosses for each *cas9* line. Efficient germline transmission of CRISPR/Cas induced mutations was observed for all *cas9* transgenes, although statistically significant differences in mean values were observed (unpaired t-test compared to *act-cas9* with the same gRNA (*, p < 0.05; **, p <0.01; ***, p <0.001)). Displayed values for *U6:3-gRNA-y* experiments are normalized to account for the 25*% y* mutant offspring that is expected in the absence of CRISPR/Cas mediated mutagenesis (Table S3). (*F*) Examples of sequences of CRISPR/Cas induced germline mutations in *e* and *y* (found in the offspring of *act-cas9 gRNA* flies). Some mutations were observed in several independent flies. The in-frame mutation in *e* (bottom sequence) was obtained from a fly with wild-type pigmentation. The ATG start codon in this and all subsequent figures is in bold.

Flies expressing the *U6:3-gRNA-e* transgene were crossed to the transgenic Cas9 lines. Remarkably, all animals expressing *gRNA-e* and *act-cas9* had mosaic pigmentation in which the majority of cuticle exhibited very dark coloration (Fig. 1*B*). This result demonstrates highly efficient biallelic targeting of *e* in somatic cells. Surprisingly, the two independent genomic insertions of *vasa-cas9* in combination with *gRNA-e* also resulted in *e* mutant tissue throughout most of the cuticle (Fig. 1*B*). Thus, Cas9 activity is not restricted to the germline by either *vasa-cas9* transgene. All *nos-cas9:GFP U6:3-gRNA-e* flies had patches of dark cuticle, but these were relatively small and infrequent (Fig. 1*B*). A similar phenotype was observed in ∼40% of flies in which *U6:3-gRNA-e* was combined with *UAS-Cas9* driven by the strong transcriptional activator Gal4VP16 under the control of *nos* regulatory sequences (*nosG4VP16*) (Fig. 1*B*). In contrast, all *nos-cas9 U6:3-gRNA-e* flies had wild-type coloration of the entire cuticle (Fig. 1*B*). The differential activity of *nos-cas9:GFP* and *nos-cas9* – which both utilize the same *nos* promoter and 3’UTR sequences and are inserted at the same genomic location – presumably reflects an influence of the GFP moiety, number of nuclear localization signals or differences in the codon usage for the Cas9 open reading frame in each construct (Table S1).

We next compared the somatic activity of the Cas9 transgenes using a second gRNA transgene, which targets the X-linked *yellow* (*y*) gene 94bp after the start codon (referred to as *gRNA-y;* Fig. 1*A*). This gRNA was also expressed from the *U6:3* promoter and was stably integrated at the *attP2* site on chromosome 3. In the absence of a wild-type *y* allele, the adult cuticle has lighter, yellow pigmentation. The combination of *U6:3-gRNA-y* with *act-cas9* or *vasa-cas9* led to a predominantly yellow cuticle in all animals (Fig. 1*C*). This was even the case in a *vasa-cas9 U6:3-gRNA-y* genotype in which four wild-type *y* alleles had to be mutated to give the yellow phenotype (Fig. 1*C*; note that different crosses had different numbers of *y*^+^ alleles due to differences in the genotypes of the Cas9 parental strains (Fig. 1*C*; Table S3; Materials and Methods)). In contrast, *nos-cas9:GFP U6:3-gRNA-y* animals had cuticle that contained only small clones of *y* mutant tissue (Fig. 1*C*), whereas the combination of *U6:3-gRNA-y* with *nosG4VP16>UAS-cas9* or *nos-cas9* resulted in phenotypically wild-type cuticle throughout the animal (Fig. 1*C*). *nos-cas9 U6:3-gRNA-y* males, in which only a single wild-type *y* allele needed to be mutated to produce yellow cuticle, also had wild-type pigmentation throughout the animal (Fig. S1). Together, the experiments with *gRNA-e* and *gRNA-y* demonstrate that *act-cas9* and *vasa-cas9* have substantial activity in cells that give rise to the cuticle, whereas activity in these cells is relatively low in *nos-cas9:GFP* and *nosG4VP16>UAS-cas9* flies and not detectable in *nos-cas9* flies.

We next assessed the germline transmission of CRISPR/Cas-induced mutations in *e* and *y* (Fig. 1*D* and *E*). In the case of *e*, this was performed by crossing individual flies containing each *cas9* transgene and *U6:3-gRNA-e* to a classical *e* mutant strain (at least eight crosses per transgene). As expected from the transgenic CRISPR/Cas approach, transmission rates were very similar for different replicates of the same cross (see error bars in Fig. 1*D*). All *cas9* transgenes resulted in 45 – 55 % of transmitted *e* alleles being non-functional (mean values per transgene) with the exception of *nos-cas9*, for which the mean value was 26±3% (Fig. 1*D*). Sequencing of non-functional *e* alleles from one of these crosses revealed a range of small insertions or deletions in the vicinity of the predicted Cas9 cleavage site (Fig. 1*F*), consistent with molecular analysis of CRISPR/Cas-induced alleles in other studies (5–8, 10).

We assessed the transmission of CRISPR/Cas-induced *y* mutant alleles by crossing flies containing *U6:3-gRNA-y* and the different *cas9* transgenes to a strain with a classical *y* mutant allele. In all cases, an average of > 70% of the progeny expected to have wild-type pigmentation in the absence of mutagenesis had phenotypically yellow cuticle (Fig. 1*E*). In the case of the *act-cas9* and one of the *vasa-cas9* transgenes, an average of 99 – 100% of such progeny were yellow (Fig. 1*E*), demonstrating CRISPR/Cas-induced mutagenesis of wild-type *y* alleles in almost every germ cell. As expected, sequencing analysis of the progeny revealed the presence of Indels at the predicted location of the *y* locus (Fig. 1*F*). Collectively our experiments indicate that *act-cas9* and *vasa-cas9* transgenes can induce mutations in somatic and germline cells with high frequencies and that *nos-cas9* allows efficient germline transmission of mutations without widespread targeting in somatic cells.

It is intriguing that for each Cas9 transgene the frequency of transmission of non-functional *y* alleles was substantially higher than that observed for *e* (Fig. 1*D* and *E*). This difference could reflect reduced target site recognition by *gRNA-e* compared to *gRNA-y*. Alternatively, in-frame Indels at the gRNA target site (which are expected to represent one third of Cas9-induced mutations) may interfere with gene function of *y* but not *e*. Compatible with this notion, a small in-frame deletion at the gRNA target site was found in one of two phenotypically wild-type progeny from *act-cas9 U6:3-gRNA-e* parents that were analyzed by sequencing (Fig. 1*F*). This finding indicates that the frequency of nucleotide changes in *gRNA-e* experiments could be higher than suggested by phenotypic screening.

### Optimized single and double gRNA expression vectors

To date, gRNAs targeting the *Drosophila* genome have been transcribed from one of the DNA polymerase III dependent promoters of the *U6* snRNA genes (5, 9–11). There are three such *U6* genes in the fly, with the vast majority of published CRISPR/Cas experiments using gRNAs produced by the *U6:2* promoter. The *U6:1* promoter has only been assessed in a single experiment using direct plasmid injection to deliver the gRNA construct (10), whereas the *U6:3* promoter has not been used previously. To enable comparison of the activity of the three *U6* promoters in CRISPR/Cas experiments, plasmids were generated harboring each *U6* promoter followed by the gRNA core sequence and a BbsI restriction site cassette (Fig. 2*A* and S2). The plasmids also contained an *attB* site to allow integration at defined positions in the *Drosophila* genome and an eye pigmentation marker to permit screening for insertion events. We used these plasmids to generate constructs expressing *gRNA-y* under the control of each *U6* promoter and stably integrated them into the *Drosophila* genome. To avoid positional effects on gRNA expression, the three *gRNA-y* transgenes were targeted to the *attP40* site on the second chromosome.

**Figure 2:**
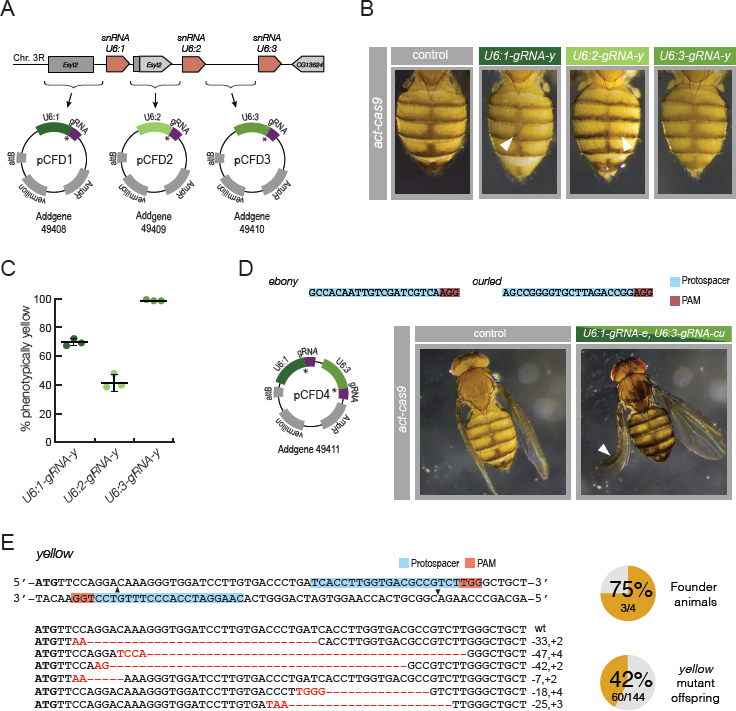
Versatile gRNA expression vectors. (*A*) Schematic of *U6* locus. Orange, *U6* genes; gray, intervening sequences and neighboring genes. Sequences 5’ to each *U6* gene were cloned in front of a core gRNA sequence to generate plasmids *pCFDl – 3*. Asterisks, positions of Bbsl cloning cassette for insertion of target sequence. (*B*) Differential activity of *U6-gRNA* constructs in epidermal cells. Female flies are shown that are heterozygous for *act-cas9* and a *gRNA-y* transgene under control of either the *U6:1, U6:2* or *U6:3* promoter. Lighter body coloration indicates CRISPR/Cas-mediated mutagenesis of the two wild-type *y* alleles present in these flies. (*C*) Germline transmission rates using different gRNA transgenes. *nos-cas9 U6-gRNA-y* females were crossed to *y* mutant males and *y* mutant offspring counted (three crosses per genotype). Displayed values are adjusted to account for the 25% *y* mutant offspring expected in the absence of mutagenesis. Large and small horizontal bars, mean and SEM. (*D*) Targeting of two genes in the same animal with a double gRNA vector, *pCFD4*. Representative images are shown of flies that express *act-cas9* in the absence or presence of a single transgene expressing gRNAs to *ebony* (*e*) and *curled* (*cu*). Arrowhead, curled wing of fly with extensive ebony pigmentation. (*E*) pCFD4 allows mutagenesis by offset-nicking in combination with *act-cas9*^*D10A*^. Top left, target sites of gRNAs close to the *y* ATG start codon (left). Arrowheads, location of nicks. Right, summary of results from injecting *pCFD4* encoding the two *y* gRNAs into transgenic *act-cas9*^*D10A*^ embryos. Crossing to *y* flies revealed that three out of four adults tested that developed from injected embryos transmitted *y* mutant alleles. Bottom left, sequence of six different Indels found in 20 yellow progeny of the founders that were analyzed.

Flies expressing each *gRNA-y* transgene were crossed to flies expressing the *act-cas9* transgene. All progeny expressing *act-cas9* and *U6:3-gRNA-y* developed a completely yellow cuticle (Fig. 2*B*). This finding indicates extremely efficient targeting of the two wild-type *y* alleles in this cross, consistent with results of combining this Cas9 transgene with the *U6:3-gRNA-y* integrated at another genomic site (Fig. 1*C*). Under the same experimental conditions, all *act-cas9 U6:1-gRNA-y* animals developed mostly yellow cuticle, but had small spots of wild-type tissue (Fig. 2*B*). In contrast, all *act-cas9 U6:2-gRNA-y* flies retained large amounts of wild-type cuticle and had many small yellow patches (Fig. 2*B*). We also produced flies expressing *nos-cas9* and each of the *U6-gRNA-y* constructs and assessed germline transmission of nonfunctional *y* alleles to the next generation as described above. Of the progeny expected to have wild-type pigmentation in the absence of CRISPR/Cas-mediated targeting, 69 ± 2%, 41 ± 3% and 99 ± 1% were phenotypically yellow when using the *U6:1, U6:2* and *U6:3* promoters, respectively (Fig. 2*C*). Collectively, these results demonstrate that the different U6 promoters differ substantially in their ability of drive mutagenesis in both somatic cells and the germline, with *U6:3* the strongest, *U6:2* the weakest and *U6:1* having intermediate activity. Simultaneous expression of two gRNAs can be used to create defined deletions (5, 8), to target two genes simultaneously (19) or for mutagenizing a single gene by offset nicking (20, 21). The latter method increases CRISPR/Cas specificity by combining a Cas9 “nickase” mutant (Cas9^D10A^ (3)), which cuts only one DNA strand, with two gRNAs to the target gene. DSBs are only induced by this protein when two molecules are guided to opposite strands of the DNA in close proximity, an event that is extremely unlikely at an off-target site. This method has not previously been used in *Drosophila*. To enable simultaneous expression of two gRNAs we produced a plasmid containing both the *U6:1* and *U6:3* promoters adjacent to gRNA core sequences (Fig. 2*D*). Different promoters were chosen to prevent the risk associated with recombination between identical sequences during cloning exercises. The plasmid was designed to allow one-step cloning of the two gRNAs by introduction of target sequences in PCR primers (Fig. S2), and therefore allows more convenient and rapid workflow than a previously reported strategy for the same task, which used sequential cloning of gRNAs adjacent to two *U6:2* promoters (8).

We used our dual gRNA plasmid to make a transgenic line simultaneously expressing gRNAs targeting *e* or *curled* (cu), which when mutated causes a curled wing phenotype (22), and crossing it to *act-cas9* flies. All 120 of the analyzed *act-cas9 gRNA-e gRNA-cu* flies had large patches of dark cuticle and 52% had one or two curled wings (Fig. 2*D*). Thus, the dual gRNA plasmid can be used to efficiently target two genes in the same animal.

We also integrated two sequences targeting opposite strands of the *y* locus into the dual gRNA vector and directly injected the plasmid into transgenic embryos expressing the Cas9^D10A^ nickase under the control of the *act* promoter (Table S1). 42% of the offspring from injected animals received non-functional *y* alleles, which we confirmed was associated with Indels at the expected position (Fig. 2*E*). All transgenic animals expressing both *act-cas9*^*D10A*^ and the strong single-target *U6:3 gRNA-y* transgene had wild-type coloration of the cuticle (data not shown), suggesting that single DNA nicks have low mutagenic potential in flies. These results demonstrate that the dual gRNA vector can be used in combination with *act-cas9*^*D10A*^ flies to create mutagenic DSBs through offset nicking.

### Efficient germline transmission of mutations in essential genes

The experiments described above identified Cas9 and gRNA reagents that can target non-essential genes in somatic and/or germline tissues with different efficiencies. We next sought to use this information to identify Cas9/gRNA combinations that can give efficient germline transmission of mutations in genes that are essential for viability. This necessitates finding conditions in which the transmission of such mutations is not compromised by deleterious effects of biallelic targeting of the gene in somatic tissues. We generated transgenic fly lines expressing gRNAs targeting the genes encoding either the secreted signaling molecule Wingless (Wg; the *Drosophila* Wnt1 ortholog) or the Wnt secretion factor Wntless (Wls). Both the *wg* and *wls* genes are essential in the soma for development of *Drosophila* to adult stages (23–25). Both gRNAs were under the control of the *U6:3* promoter and integrated at the *attP40* site. *gRNA-wg* and *gRNA-wls* direct Cas9 to cut, respectively, just after or within the region of the target gene that encodes the signal sequence (Fig. 3*A*).

**Figure 3:**
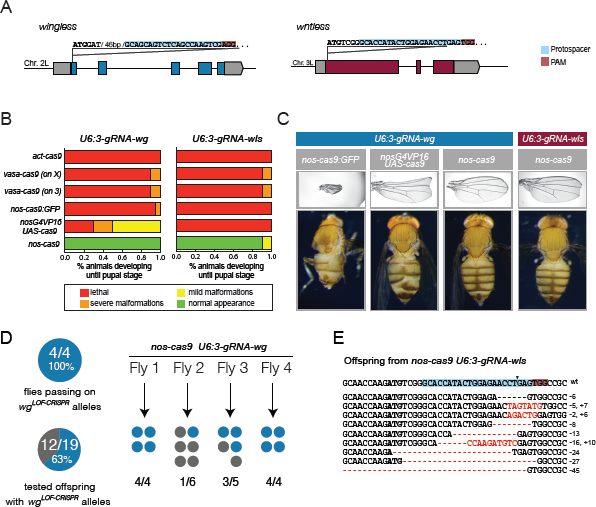
Efficient targeting of essential genes. (*A*) Schematic of sequence and position of target sites of *gRNA-wg* and *gRNA-wls*. (*B*) Summary of fate of animals that developed to pupal stages that co-expressed *U6:3-gRNA-wg* or *U6:3-gRNA-wls* with different *cas9* transgenes. (*C*) Representative examples of phenotypes observed in adult *cas9 gRNA-wg* and *cas9 gRNA-wls* flies. Defects consistent with perturbed Wg signaling include wing notches (upper panels), and malformation of the legs and abdominal segments (lower panels)(left and center left panels exemplify severe malformation; right panel exemplifies mild malformation). (*D*) Assessment of transmission of non-functional *wg* alleles. *nos-cas9 gRNA-wg* flies were crossed to a balancer fly line. 19 flies from 4 different parents were then tested for transmission of non-functional *wg* alleles by genetic complementation with a known *wg* null allele. 12 flies received *wg* loss-of function alleles (*wg*^*LOF-CRISPR*^) that failed to complement *wg*^−^. Note that *wg*^*CRISPR*^ alleles that retain function are not detected by this method. (*E*) Assessment of transmission of nonfunctional *wls* alleles. A fertile *nos-cas9 gRNA-wls* fly was crossed to a balancer stock and the *wls* locus of 15 heterozygous progeny sequenced. 10 sequencing chromatograms could be unambiguously read and all contained Indels at the target site. The mutated sequence is shown below the wild-type sequence. The 27bp deletion was recovered in two independent flies.

When *U6:3-gRNA-wg* or *U6:3-gRNA-wls* flies were crossed to the ubiquitous *act-cas9* line, no offspring survived past the pupal stage (Fig. 3*B*), presumably due to effective biallelic targeting of *wg* and *wls* in the soma. Crossing *vasa-cas9, nos-cas9:GFP* and *nosG4VP16>UAS-cas9* to each gRNA also resulted in substantial lethality, with the few animals that progressed to adulthood exhibiting a range of morphological defects consistent with defects in Wg signaling (Fig. 3*C*). Unsurprisingly, many of these adults died shortly after eclosion and those that survived often had poor fertility. In contrast, no significant lethality was observed for *nos-cas9 U6:3-gRNA-wg* and *nos-cas9 U6:3-gRNA-wls* pupae (Fig. 3*B*). All of the eclosed adults appeared morphologically normal, with the exception of 5% of *nos-cas9 U6:3-gRNA-wls* flies that had small notches in the wing (Fig. 3*C*). This phenotype is indicative of a local impairment of Wg signaling, thereby providing evidence that not all Cas9 activity is restricted to the germline when *nos-cas9* is combined with this gRNA. This series of experiments demonstrate that only our *nos-cas9* line was sufficiently germline restricted to allow efficient development of flies to adulthood when the essential genes *wg* and *wls* were targeted.

We next evaluated germline transmission of CRISPR/Cas induced mutations in *wg* and *wls* using *nos-cas9*. We randomly selected 19 offspring of *nos-cas9 U6:3-gRNA-wg* flies and tested for *wg* mutations by genetic complementation with a previously characterized *wg* null allele (26). In 12 of the 19 crosses the *wg* null allele was not complemented, strongly indicating the creation of non-functional *wg* mutations by CRISPR/Cas (Fig. 3*D*). Indeed, sequencing the *wg* locus confirmed Indels at the gRNA target site in all 12 flies tested (data not shown). The high percentage of germline transmission of non-functional *wg* alleles from *nos-cas9 U6:3-gRNA-wg* flies indicated that it would have been feasible to recover novel mutations in this essential gene without the need to phenotypically screen offspring prior to sequencing.

We discovered that *nos-cas9 U6:3-gRNA-wls* females were poorly fertile, which is compatible with the requirement for maternal Wls during embryonic development (24, 25). *nos-cas9 U6:3-gRNA-wls* males also had low fertility, raising the possibility of a novel role for Wls in the male germline. Nonetheless, we were able to sequence *wls* alleles from PCR products of 15 randomly selected offspring of an outcrossed *nos-cas9 U6:3-gRNA-wls* fly that was fertile. The sequencing chromatograms had double peaks around the gRNA target site in every case. This pattern indicates the presence of a CRISPR/Cas-induced *wls* mutant allele in each fly, along with the wild-type allele inherited from the other parent. Peaks from the mutagenized and wild-type *wls* alleles could be called unambiguously in the sequencing traces from 10 of the 15 samples; these 10 traces revealed 10 Indel mutations, nine of which were unique (Fig. 3*E*). Many of the Indels created frameshifts or removed the ATG start codon, suggesting that they were null mutations. Together these results show that our optimized CRISPR/Cas system allows the generation of loss-of function alleles in essential genes at frequencies that make screening with methods other than DNA sequencing unnecessary.

### Precise genome modification by homologous recombination

The work described above revealed tools to generate loss-of-function Indel mutations in both the soma and germline of *Drosophila*. In many cases, it is also desirable to further dissect gene function by introducing specific sequence changes into the genome. In *Drosophila* it has been possible for several years to create specific genome modifications by homologous recombination, although targeting typically occurs with very low efficiency (27). Recent studies have suggested that the CRISPR/Cas system can increase the efficiency of homologous recombination in flies, implying that the generation of site-specific DSBs is rate limiting for homologous recombination (11, 26). However, in these studies the frequency of precise integrations by HR was still sufficiently low that flies harboring the desired integration had to be identified by a linked visible marker. We therefore tested whether the highly effective creation of DSBs by our transgenic CRISPR/Cas system could be exploited to facilitate efficient homologous recombination.

We designed an oligonucleotide donor to introduce an 11-bp sequence containing a recognition sequence for the BglII restriction enzyme at the *gRNA-e* target site in the *e* locus and injected the donor DNA into *act-cas9 U6:3-gRNA-e* embryos (Fig 4*A*). 76 progeny of five flies that developed from the injected embryos were selected at random (i.e. without consideration for their pigmentation phenotype) and a diagnostic BglII digest performed on PCR products amplified from their *e* locus (Fig. 4*B*). Three of the five tested flies were found to have transmitted the designed mutation, passing it on to 7 – 40% of their offspring (Fig. 4*C*). In total, 8 of the 76 tested progeny (11%) carried the mutation (Fig. 4*C*). Sequencing of the genomic locus confirmed the precise integration of the donor DNA (Fig. 4*D*). These data demonstrate that our CRISPR/Cas tools can allow homologous recombination of oligonucleotide-derived sequences with efficiencies compatible with directly screening for targeting events by PCR and sequencing.

**Figure 4:**
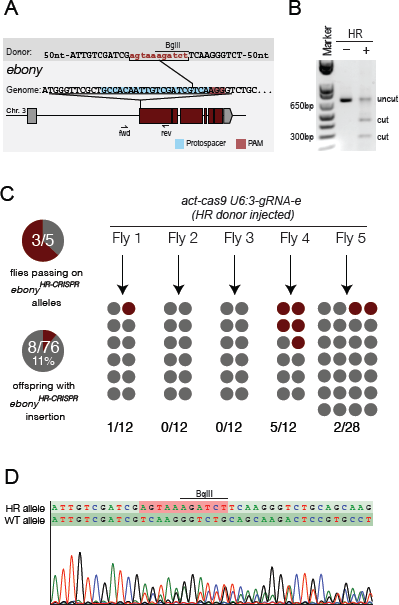
Efficient incorporation of an exogenous sequence by homologous recombination following Cas9-induced DSBs. (*A*) Schematic of the donor DNA in relation to the gRNA target site at the *e* locus. The donor was a single stranded oligonucleotide with 60-nt homology to the target locus at either side of the Cas9 cut site and an 11-nt insert (lower case). This insert introduces an in-frame stop codon (TAA) and a BglII restriction site. The locations of the primers used for the genotyping PCR in *B* are shown below the schematic of the genomic locus. Donor DNA was injected into embryos that were the progeny of *act-cas9* females and *U6:3-gRNA-e* males. (*B*) Successful integration of the donor construct could be detected in the offspring of injected embryos by BglII digestion of PCR products. Agarose gel showing pattern observed in the absence (–) and presence (+) of the homologous recombination (HR) event. The 700-nt fragment present in both samples is derived from the wild-type *e* locus transmitted by the other parent. (*C*) Summary of results from screening flies for HR events by PCR and restriction digest. (*D*) Sequence verification of the precise integration of the donor DNA in the *e* locus by direct sequencing of a PCR product amplified from a heterozygous fly that tested positive by BglII restriction digest. Double peaks in the chromatogram represent an overlay of the sequence of the mutant and wild-type *e* alleles.

### CRISPR/Cas as a tool for analyzing loss-of-function mutant phenotypes in somatic tissues

An important result from the evaluation of our tools is that transgenic CRISPR/Cas with *U6:3-gRNAs* can result in very effective gene targeting in the soma, including mutagenesis of multiple alleles of the same gene. We noticed in our experiments targeting *y* and *e* that cuticles frequently contained large patches of cells with the same mutant phenotype (Fig. 1*B* and *C*), which presumably represent clonally derived cells carrying the same CRISPR/Cas-induced lesion. Analysis of classical *Drosophila* mutations in genetic mosaics has been invaluable for understanding functions of genes that act at multiple stages during development, as well as cell autonomous and non-autonomous mechanisms (28). However, generating the stocks required to induce clones by traditional methods is time consuming (see Discussion). We therefore wondered if our CRISPR/Cas tools could be used to rapidly generate clones that have both alleles of essential genes mutated.

We first analyzed wing imaginal discs from *act-cas9 U6:3-gRNA-wls* third instar larva using an antibody that recognizes the C-terminal tail of the Wls protein (29). Endogenous Wls is expressed ubiquitously in the wing imaginal disc, where it is required for the secretion of the Wg signaling molecule from Wg producing cells (24, 25, 29). All wing imaginal discs from control *act-cas9 U6:3-gRNA-e* animals had wild-type Wls protein levels and distribution (Fig. 5*A*). In contrast, all *act-cas9 U6:3-gRNA-wls* wing imaginal discs examined contained patches of cells with reduced Wls levels, no Wls protein and wild-type Wls levels (Fig. 5*B*). These first two scenarios presumably reflect clonal tissue that has, respectively, one or two *wls* mutant alleles that fail to produce full length Wls protein. Wg protein was retained in Wg producing cells within clones that lacked detectable Wls protein (Fig. 5*B*), confirming biallelic disruption of *wls*.

**Figure 5:**
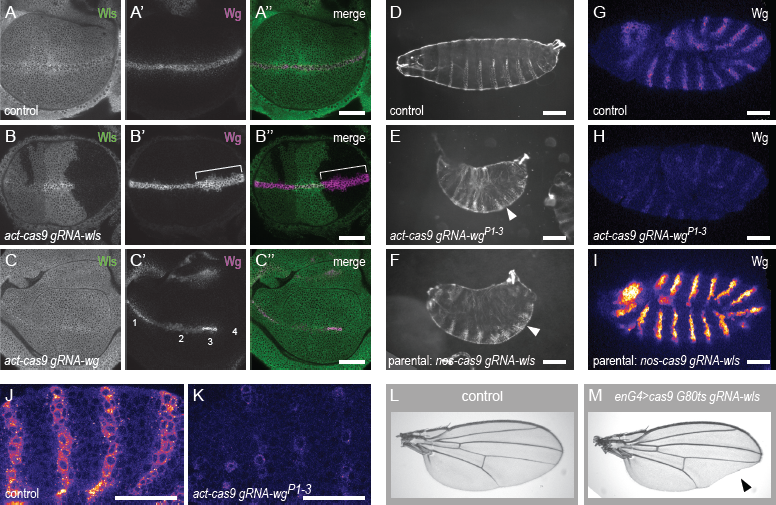
Revealing mutant phenotypes through efficient biallelic targeting with transgenic CRISPR/Cas. (*A – C*) Genetic mosaics induced in third instar wing imaginal discs by somatic CRISPR/Cas. (*A*) Control *act-cas9 U6:3-gRNA-e* wing discs have ubiquitous expression of Wls and Wg expressed in a stripe along the dorsal-ventral boundary. (*B*) *act-cas9 U6:3-gRNA-wls* discs have patches of cells with wild-type Wls levels, reduced Wls expression and no detectable Wls protein. Wg protein accumulates intracellularly in Wg producing cells in the absence of Wls (bracket). (*C*) Clones with different distributions of endogenous Wg protein can be observed in *act-cas9 U6:3-gRNA-wg* discs: (1) wild-type, (2) nuclear accumulation (3), increased Wg levels and (4) no Wg protein. A second example with abnormal nuclear Wg staining is shown in higher magnification in Fig. S3. Images are representative of >10 discs of each genotype. (*D – K*) Biallelic targeting by transgenic CRISPR/Cas can reveal mutant phenotypes during embryogenesis. (*D – F*) Cuticle preparations at the end of embryogenesis. (*D*) *act-cas9 U6:3-gRNA-e* control animal. (*E, F*) All *act-cas9 gRNA-wg*^*P1-3*^ embryos (*E*) and most embryos from *nos-cas9 U6:3-gRNA-wls* parents (*F*) have naked region of cuticle replaced with denticle bands (e.g. arrowheads). (*G – K*) Analysis of Wg protein in stage 9/10 embryos by immunostaining (“fire” lookup table reveals full dynamic range of Wg levels in different genotypes). Compared to control *nos-cas9 U6:3-gRNA-e* embryos (*G*), Wg levels are strongly reduced in *act-cas9 gRNA-wg*^1–3^ embryos (*H*) and strongly increased in embryos from *nos-cas9 U6:3-gRNA-wls* parents (*I*). (*J, K*) Higher magnification views of Wg protein in embryos, showing variation in Wg levels in *act-cas9 gRNA-wg*^*p1–3*^ embryos (*K*), presumably due to independent CRISP/Cas targeting events in subsets of cells. *act-cas9 U6:3-gRNA-e* is shown as a control (*J*). Images are representative of > 50 embryos examined. (*L, M*) Cas9 activity can be focused on specific tissues by Gal4/UAS. Expression of *gRNA-wls* together with *UAS-Cas9* (line CFD5 (Table S1)) under the control of *enGal4* and *Gal80*^*ts*^ gives rise to viable flies that have wing notches posteriorly in the adult wing (*M*; arrowhead). Control wing (*L*) is from an animal that did not inherit the *gRNA-wls* transgene. All genotypes in this figure refer to one copy of each transgene. Scale bars represent 40 μm in *A – C*; 100 μm in *D – F*; 30 μm in *G – K*.

We next examined wing imaginal discs from *act-cas9 U6:3-gRNA-wg* animals using an anti-Wg antibody (30). The Wg expression domain normally includes a stripe of cells along the dorsal-ventral boundary of the wing pouch, a pattern always observed in discs from control *act-cas9 gRNA-e* larva (Fig. 5*A*). In contrast, *act-cas9 gRNA-wg* wing imaginal discs had disrupted patterns of Wg expression (Fig. 5*C*). Although clones that lost all detectable Wg protein were common, presumably reflecting biallelic loss-of-function mutations, we additionally observed clones with increased abundance or altered subcellular distribution of the protein (Fig. 5*C* and S3). These latter clones likely contain in-frame mutations at the gRNA target site that confer new properties to the Wg protein. This finding suggests that CRISPR/Cas can also be used to shed light on the functional requirements for specific regions of a protein.

The experiments described above demonstrated that *act-cas9* and *U6:3-gRNA* can be used to stochastically induce clones in imaginal discs that have biallelic targeting of essential genes. To investigate whether our transgenic CRISPR/Cas tools can reveal somatic mutant phenotypes at earlier developmental stages, we analyzed embryos expressing both *act-cas9* and *U6:3-gRNA-wg. wg* is expressed zygotically in the embryo and is required for segment polarity; classical *wg* null mutants fail to complete embryogenesis and have the replacement of regions of naked cuticle with an array of denticles (23). Our earlier finding that many *act-cas9 U6:3-gRNA-wg* embryos develop to pupal stages demonstrates that they retain significant *wg* function (Fig. 3*B*). We therefore generated a new gRNA transgene that simultaneously expressed three gRNAs targeting the *wg* promoter (*gRNA-wg*^*p1–3*^). This construct, in which the gRNAs were under the control of the *U6:1, U6:2* and *U6:3* promoters, was integrated at the *attP2* site. All *act-cas9 gRNA-wg*^*P1-3*^ animals arrested before the end of embryogenesis with a strong segment polarity phenotype (Fig. 5*D* and *E*). This result presumably reflects biallelic disruption of *wg* expression because one functional copy of *wg* is sufficient for normal development. Indeed, immunostaining of *act-cas9 gRNA-wg*^*P1-3*^ embryos revealed strongly reduced expression of Wg protein compared to *act-cas9 U6:3-gRNA-e* controls at all stages examined (Fig. 5*G, H, J, K*), indicative of interference with transcription of both alleles. We conclude that somatic CRISPR/Cas can be used to induce biallelic mutations in genes that are expressed zygotically in embryos.

Additional observations indicated that transgenic CRISPR/Cas can also be used to study maternal gene function in the embryo. As described above, *nos-cas9 U6:3-gRNA-wls* adults had low fertility. Closer inspection revealed that intercrosses of these flies resulted in many eggs being laid, but with less than 5% developing to larval stages. Wls is normally contributed maternally to the embryo. Whereas zygotic *wls* null mutants from heterozygous mothers develop until pupal stages, offspring of mothers with a homozygous *wls* mutant germline die at embryonic stages with cuticle phenotypes typical of perturbed Wg signaling (24, 25). The vast majority of embryos from *nos-cas9 U6:3-gRNA-wls* mothers arrested at the end of embryogenesis with segment polarity phenotypes reminiscent of *wls* germline mutant clones (Fig. 5*F*). This observation indicates biallelic disruption of *wls* in the germline of *nos-cas9 U6:3-gRNA-wls* females. Consistent with this notion, immunostaining revealed a strong accumulation of Wg protein in producing cells in embryos derived from *nos-cas9 U6:3-gRNA-wls* mothers (Fig. *5I*), which is presumably due to defective Wg protein secretion in the absence of Wls. These findings reveal that CRISPR/Cas can also be used to study loss-of-function embryonic phenotypes of maternally contributed genes.

Finally, we explored whether biallelic gene targeting in specific somatic cell types was possible by coupling CRISPR/Cas mediated mutagenesis to the UAS-Gal4 system. *UAS-cas9* was combined with *engrailed-Gal4* (*enGal4*), which drives *UAS* expression specifically in posterior compartments of the animal, and *gRNA-wls*. We also included a temperature sensitive Gal80 transgene (*Gal80*^*ts*^) to restrict Cas9 activity further by temporal control of Gal4. Following a temperature shift of second instar larvae of this genotype to 29°C, many adult flies had pronounced notches specifically in the posterior half of the wing (Fig. *5L* and *M*). This phenotype was not observed in the absence of *gRNA-wls* (Fig. *5L*). These results show that biallelic targeting of essential genes can be focused on specific cell types by regulating expression of Cas9 with UAS-Gal4. However, the first generation of *UAS-cas9* constructs requires optimization. Substantial lethality was observed when *UAS-cas9* transgenic lines were crossed to strong, ubiquitous GAL4 drivers, which was not dependent on the presence of a gRNA. This toxicity was also not dependent on the endonuclease activity of Cas9 as it was also observed with a catalytically dead mutant protein. Furthermore, when *UAS-cas9* was combined with *gRNA-wls* we observed some *wls*^−^ clones in wing discs that were independent of Gal4 activity (data not shown). Nonetheless this effect, which is presumably due to some leaky expression from the *UAS* promoter, was not sufficient to produced wing notching in the adult. Future efforts will be aimed at producing *UAS-cas9* variants that have reduced expression levels yet still retain sufficient activity for biallelic targeting in response to Gal4.

## Discussion

Genome engineering with CRISPR/Cas promises to revolutionalize genome engineering in a large variety of organisms. Whereas proof-of-principle studies have demonstrated CRISPR/Cas-mediated mutagenesis in *Drosophila melanogaster* (5–11), a systematic evaluation of different tools has not been reported. We have developed a collection of tools to further harness the potential of CRISPR/Cas for functional studies in *Drosophila*. We have characterized different Cas9 and gRNAs expression constructs and revealed combinations that allow mutagenesis of essential and non-essential genes with high rates, support efficient integration of exogenous sequences and can readily reveal mutant phenotypes in somatic tissues.

To assess the activity of different constructs we used a fully transgenic CRISPR/Cas system in which Cas9 and gRNA are expressed from plasmid sequences stably integrated at defined positions in the genome. The advantage of this approach for comparative analysis is the low variability between experiments compared to protocols that use direct microinjection of RNA or DNA. For example, we found that 100% of flies expressing transgenic Cas9 and gRNA constructs had efficient germline mutagenesis of the target site. In contrast, the number of animals that transmit mutations to the next generation using microinjection-based delivery of one or both CRISPR/Cas components is highly variable and often less than 50% (5, 6, 9–11).

Our quantitative analysis using transgenic Cas9 and gRNA constructs reveals that, of all three *U6* promoters, the previously uncharacterized *U6:3* promoter leads to the strongest gRNA activity. This observation suggests that targeting rates of previous approaches have been limited by the selection of the *U6:2* or *U6:1* promoter. In combination with highly active Cas9 lines (e.g. *act-Cas9, vasa-cas9*), transgenic *U6:3-gRNA* can be used to transmit mutations in non-essential genes through the germline with remarkably high efficiency (45 – 100% of offspring receiving a non-functional CRISPR-generated allele). However, these Cas9 lines are also active outside the germline and biallelic targeting in the soma effectively prevents their use for transmission of essential genes in combination with *U6:3-gRNA* transgenes. The ubiquitous activity of *vasa-cas9* was presumably masked in a previous study in which a *U6:2-gRNA* plasmid targeting the eye pigmentation gene *rosy* was injected at the posterior of the embryo, thereby restricting its activity in the vicinity of the future germ cells (11). It remains to be evaluated if activity of CRISPR/Cas is sufficiently restricted by this approach to allow efficient germline transmission of mutations in essential genes. We demonstrate that efficient targeting of essential genes is possible using *U6:3-gRNA* transgenes in combination with the transgenic *nos-cas9* line in which endonuclease activity is largely germline restricted. This was even the case for *wls*, which is required in the germline, because fertility (although much reduced) was not abolished. It is conceivable that efficient targeting of genes that are cell lethal will necessitate further reduction of endonuclease activity to avoid biallelic disruption in the germ cells. This could be achieved by using transgenic *U6:2-gRNA* constructs or delivering a dilution series of a *U6:3-gRNA* plasmid by embryo injection.

Our optimized CRISPR/Cas system also allowed the integration of a small designer sequence with high efficiency. By injecting an oligonucleotide donor into embryos expressing *cas9* and *gRNA* transgenes we achieved integration at the target site in 11% of all offspring analyzed. A previous study in which a similarly sized donor construct was injected together with *cas9* and *gRNA* plasmids found that 0.3% of all offspring integrated the exogenous sequence at the target site ((5) and S. Gratz personal communication). Very recently, Gratz et al. have reported integration rates for larger constructs that range from 0 – 11% by injection of the donor and gRNA plasmid into *vasa-cas9* transgenic embryos (11). It is difficult to directly compare the integration efficiency with our method to the previous studies due to the use of different gRNAs and donors. Nonetheless, the increased reproducibility and efficiency of site-specific DSB generation with our optimized fully transgenic CRISPR/Cas is likely to facilitate integration of exogenous sequences by homologous recombination compared to methods that deliver Cas9 or gRNA by direct embryo injection. The frequency of integration is of paramount importance when the goal is to introduce subtle changes in the DNA sequence, such as individual point mutations. This is because screening for rare insertion events by co-integration of a selectable or visible marker gene is problematic due to the retention of exogenous sequences that can affect function of the targeted locus (note that exogenous sequences are still retained following removal of markers by site-specific recombinases (31)). We demonstrate here that optimized transgenic CRISPR/Cas can facilitate the introduction of precise changes at endogenous loci with rates that make it practical to screen solely by PCR and Sanger sequencing. We envisage that one important application of our tools will be the introduction of disease mutations into endogenous loci within the *Drosophila* genome in order to model human pathologies.

We also demonstrate a novel application of CRISPR/Cas in rapidly analyzing gene function in somatic cells. Our optimized tools permit efficient generation of genetic mosaics in which clones of cells have biallelic gene disruption. Mosaic analysis is a powerful method to study genes that control cell interactions in complex tissues or function at multiple stages of development (28). Mitotic recombination using the yeast site-specific recombinase Flp is typically used for this purpose in *Drosophila* (31). This method requires recombination of the mutation of interest onto a chromosome containing the FRT recognition sites for Flp, which is a time-consuming process (particularly for genes located close to the FRT site). Somatic CRISPR/Cas requires only one cross to give rise to mosaics. Furthermore, unlike the FRT-Flp system, somatic CRISPR/Cas is not dependent on cell division and hence should be better suited to interrogate gene function in differentiated tissues such as the adult brain. One advantage of FRT-Flp for mosaic analysis is that clones can be readily marked by the presence or absence of a linked marker gene, such as *gfp*. Here we identified mutant cells with antibodies specific to the proteins studied but for many proteins no antibody will be available. Under these circumstances we envisage mutant tissue being identified by first using CRISPR/Cas to generate a line in which a fluorescent protein or short epitope tag has been knocked in to the endogenous locus. Such a reagent would in any case be valuable for characterizing function of the protein. In the future, CRISPR/Cas and FRT-Flp will probably co-exist as complementary methods for mosaic analysis of gene function.

As is the case for somatic CRISPR/Cas, RNAi can reveal mutant phenotypes after a single cross. However, RNAi often reduces gene expression only partially, meaning that some mutant phenotypes can be incompletely penetrant or missed all together due to residual protein. Somatic CRISPR/Cas readily provides access to null mutant phenotypes caused by biallelic out-of-frame mutations. Consequently, somatic targeting of all genes tested (*e*, *y*, *cu, wg* or *wls*) could reveal the classical loss-of-function mutant phenotype, and with high penetrance. Furthermore, RNAi is notorious for off-target effects (32), which can complicate phenotypic analysis significantly. Early studies in human cells did suggest that off-target mutagenesis is also a prevalent problem of CRISPR/Cas (33, 34). However, experiments in *Drosophila* have so far failed to detect induced mutations at sites without perfect complementarity (6, 11). These findings suggest that CRISPR/Cas operates with high fidelity in flies, although addressing the issue of specificity definitively will require genome wide analysis of DSB induction. Nonetheless, it is encouraging that nonspecific phenotypes were not detected in any of our experiments targeting e, y, *cu, wg* or wls. A recent study showed that truncating the target sequence of gRNAs results in improved fidelity of CRISPR/Cas in human cells (35), suggesting a straightforward approach to minimize the risk of off-target effects in *Drosophila*. Our generation of tools for offset nicking-based mutagenesis in flies provides a further option for increasing specificity. Such is the ease with which CRISPR/Cas experiments can be performed, the specificity of mutant phenotypes can also be tested using multiple independent gRNAs.

High throughput genetic screens are likely to be another important application of somatic gene targeting by CRISPR/Cas. The generation of gRNA expression plasmids through cloning of annealed short oligonucleotides is an easy and economic one-step process that is scalable to generate a library of targeting vectors for thousands of genes. CRISPR/Cas based screens will be particularly powerful if mutagenesis can be restricted in time and space. Here we take a step in this direction by demonstrating that expressing Cas9 under the control of the Gal4/UAS system can enrich biallelic targeting within a defined group of cells.

CRISPR/Cas genome engineering is rapidly changing basic and applied research. We hope that the results described in this study will further the development of genome engineering in *Drosophila* and contribute to the rapid incorporation of this technique into the toolbox of every fly geneticist.

## Material and Methods

Addition Materials and Methods are described in the *Supplementary Material and Methods*.

### Plasmid construction

Unless otherwise noted, cloning was performed by Gibson assembly (using Gibson assembly master mix, New England Biolabs). PCR products were produced with the Q5 2x mastermix kit (NEB). All inserts were verified by sequencing. Details on plasmid construction are available in the *Supplementary Material and Methods*. Primers used for plasmid construction are listed in Table S4.

#### *Drosophila* genetics

Details of transgenes and fly stocks are given in Tables S1 – 3 and S5. Throughout this study, transgenic *cas9* virgin females were crossed to *U6-gRNA* expressing males.

##### Assessing relative activity of Cas9 and gRNA lines

*cas9* virgin females were crossed to transgenic gRNA males. The genotype of the individual crosses is given in Table S3. All crosses involving *gRNA-e* produced offspring with two wild-type *e* alleles that can be subject to gene targeting by CRISPR/Cas. Since some Cas9 strains were generated in a *y* mutant background and some attP sites are marked by *y+* transgenes, offspring from the different *gRNA-y* crosses inherit different numbers of *y* alleles from their parents (Fig. 1 and Table S3). As a result, either 0% or 25% of F2 offspring from the different crosses would be expected to be phenotypically yellow in the absence of CRISPR/Cas mediated mutagenesis. To correct for this imbalance we subtracted 25% of the phenotypically yellow animals when calculating germline transmission rates of non-functional alleles from data collected from the latter type of cross.

##### Genetic complementation of wg loss-of-function alleles

*nos-cas9/+; gRNA-wg/+* flies were crossed to a *Sp/CyO* balancer strain. Individual flies of the genotype *wg*^*test*^/*CyO* were then crossed to *wg*^*TV-Cherry*^/*CyO* flies. *wg*^*TV-Cherry*^ is a *wg* null allele in which the first exon of *wg* is replaced with a homologous recombination targeting cassette (26). The next generation of the crosses was then screened for the presence or absence of flies without the *CyO* chromosome. Cultures in which all flies had the *CyO* chromosome indicated that the *wg*^*test*^ allele could not genetically complement *wg*^*TV-Cherry*^ and hence did not encode functional Wg protein.

### Molecular characterization of target loci

To identify the molecular nature of CRISPR/Cas induced mutations genomic DNA was extracted from individual flies by crushing them in 15 μl microLysis-plus (Microzone, UK) and releasing the DNA in a thermocycler according to the supplier’s instructions. 0.5 μl of the supernatant was used in 25 μl PCR reactions (Q5 2x mastermix kit (NEB)) using primers binding 300 – 500 bp upstream and downstream of the target site. PCR products were gel purified and either directly sent for Sanger sequencing or cloned into pBluescript-SK(+) by Gibson assembly. In the latter case, single colonies were selected as templates for PCR amplification and PCR products sequenced. Direct sequencing of PCR products from genomic DNA for heterozygous flies usually yields an overlay of the DNA sequence from both chromosomes. Indels can be observed as regions with double peaks in the trace. In the majority of such regions the sequence of both alleles can be unambiguously called (see Fig. 4*D* for an example).

### Embryo injections

Embryos were injected using standard procedures (further details are given in the *Supplementary Materials and Methods*). For the delivery of plasmid DNA for the production of transgenes with the Phi31C system, 150ng/μl of DNA in sterile dH_2_O was injected. Single stranded DNA oligonucleotides designed to modify the *e* locus were injected into the posterior region of *act-cas9/+; U6-3-gRNA-e/+* embryos as a 500 ng/μl solution in dH_2_O.

## Acknowledgements

We would like to thank Cyrille Alexandre, Alberto Baena-Lopez, Kelly Beumer, Dana Carroll, Scott Gratz, Melissa Harrison, Kate Koles, Kate O’Connor-Giles, Avi Rodal, Jean-Paul Vincent, Jill Wildonger, and others from the *Drosophila* genome engineering community for generously sharing unpublished results and reagents. We are also very grateful to Madalena Reimao Pinto and Tamsin Samuels for help with cloning and genotyping and Silvia Aldaz, Konrad Basler, Johannes Bischof, Simon Collier, Justin Crooker, Norbert Perrimon, Katja Röper, David Stern and Ryohei Yagi for additional reagents. This study was supported by a Marie-Curie Intraeuropean Fellowship (to F.P.), Howard Hughes Medical Institute (H.-M.C and T.L) and the UK Medical Research Council (S.L.B.; Project number U105178790).

## Author contributions

F.P. conceived study; F.P. and S.L.B designed experiments; F.P. performed experiments; F.P. and S.L.B. analyzed data; F.P. and S.L.B. wrote the manuscript; H.-M.C. and T.L. first discovered the difference in activity between *U6:2* and *U6:3* promoters and contributed one of the *UAS-cas9* lines.

